# Instrument-specific learning improves *de novo* peptide sequencing from timsTOF spectra

**DOI:** 10.64898/2026.11.19.767847

**Authors:** Michael Dammann, Bhavya Krishnan, Sander Willems, Justin Sanders, Fabian J. Theis, Wout Bittremieux, George Rosenberger, William Stafford Noble

## Abstract

Deep learning has substantially advanced *de novo* peptide sequencing, yet most models have been developed using Orbitrap mass spectra, limiting their ability to recover peptide sequence information from other mass spectrometry platforms. Here, we demonstrate how instrument-specific learning can overcome this limitation by introducing Casanovo_tims_, a transformer-based model trained on nearly seven million peptide–spectrum matches from trapped ion mobility–time-of-flight (timsTOF) mass spectrometry data. On a held-out timsTOF benchmark, Casanovo_tims_ substantially outperforms existing Orbitrap-trained *de novo* sequencing methods, achieving over fourfold higher peptide prediction coverage at 90% precision and more than eightfold higher coverage at 95% precision compared with the strongest baseline. Importantly, these gains extend beyond benchmark performance to challenging biological applications. In immunopeptidomics, high-confidence Casanovo_tims_ predictions increase the number of unique peptides recovered from an HLA class I pulldown by 15% over database search alone, including 280 non-canonical candidates absent from the reference proteome. In gut metaproteomics, Casanovo_tims_ adds 16% more peptides confirmed in a 13.8-million-cluster microbial catalogue and proposes further plausible sequences that are not included in the catalogue. Together, these findings demonstrate that instrument-specific learning can recover peptide sequence information missed by existing *de novo* sequencing models, expanding opportunities for peptide discovery in complex biological samples where reference-based detection remains incomplete.

## Introduction

High-resolution tandem mass spectrometry (MS/MS) has enabled comprehensive proteome characterization across increasingly challenging experimental designs [1]. When high-quality sequence databases or spectral libraries are unavailable, for example in studies of non-model organisms [2], metaproteomics [3], immunopeptidomics [4], or sequence-variant discovery [5], peptide detection via database search is challenging, and *de novo* peptide sequencing provides a complementary solution. Recent deep learning approaches have substantially advanced the accuracy of *de novo* sequencing, outperforming classical algorithmic methods [6].

Among these approaches, transformer-based architectures have proven particularly effective for modeling fragment ion patterns and capturing long-range dependencies in peptide sequences. Casanovo, which was the first transformer-based *de novo* peptide sequencing tool [7], remains one of the leading open-source solutions and has been adopted in diverse workflows and development of new architectures [8–13]. However, like most contemporary models, Casanovo has been trained almost exclusively on Orbitrap higher-energy collisional dissociation (HCD) data, primarily from widely used resources such as MassIVE-KB [14] and ProteomeTools [15].

In parallel, Bruker timsTOF instruments, which couple quadrupole time-of-flight (Q-TOF) analysis with trapped ion mobility spectrometry (TIMS), have seen rapid uptake in proteomics. Although both Orbitrap HCD and Q-TOF collision-induced dissociation employ beam-type fragmentation, differences in mass analyzers, acquisition settings, and signal processing produce distinct spectral characteristics [16]. These differences have motivated instrument-specific models for fragment-ion intensity prediction, such as Prosit [17] and MS2PIP [18], while finetuning on timsTOF data has been shown to improve peptide–spectrum match (PSM) rescoring [16, 19, 20]. Consequently, *de novo* sequencing models trained predominantly on Orbitrap spectra may not generalize well to timsTOF data.

An additional consideration is the ion mobility dimension introduced by TIMS [21]. In data-dependent acquisition–parallel accumulation–serial fragmentation (ddaPASEF), ions are separated by mobility before targeted fragmentation, providing an additional dimension alongside *m/z* and retention time (RT). Although ion mobility separation can improve precursor selectivity and signal-to-noise, it also complicates representation and learning: a single MS/MS event may be associated with multiple ion mobility frames or require aggregating peaks across a mobility window.

To address these challenges, we have developed a timsTOF-native version of Casanovo, which we termed Casanovo_tims_. Our contributions are fourfold. First, we curate a large training dataset (millions of PSMs from thousands of runs) to enable competitive Casanovo_tims_ performance. Second, we introduce a data pipeline that natively ingests raw timsTOF data and internally collapses the ion mobility dimension to reduce spectrum complexity, leveraging the existing Python mass spectrometry ecosystem. Third, we evaluate Casanovo_tims_ on PSMs from peptides never seen during training as well as two challenging downstream applications, low-input immunopeptidomics and metaproteomics, demonstrating strong performance in all three settings. Fourth, we release this capability in Casanovo v5.2.2, adding a frequently requested feature for users working on timsTOF proteomics, and, to our knowledge, representing the first open-source deep learning-based *de novo* peptide sequencing model with native support for timsTOF data.

Collectively, this work expands *de novo* peptide sequencing to a widely used instrument class. By handling ion mobility natively, the new Casanovo version broadens applicability to high-throughput and low-input settings where timsTOF instruments excel.

## Results

### timsTOF and Orbitrap spectra exhibit distinct characteristics

To characterize the spectral differences between timsTOF and Orbitrap spectra relevant to *de novo* peptide sequencing, we compared MS/MS spectra of identical peptide precursors acquired on timsTOF and Orbitrap instruments. We first examined a representative pair of spectra that were generated on different instrument models but, according to database search, were generated from the same underlying peptide. The resulting mirror plot (Fig. 1a) suggests two conclusions. First, the timsTOF spectrum is more complex, with many low and medium intensity peaks. Applying a naive noise threshold to remove these peaks is not feasible because it would also remove true fragment peaks. In this example, peaks such as y_2_, y_5_, and b_10_ would likely not survive such a filter. Second, the relative intensities of the fragment ions differ between the two instruments, with most peaks only appearing in one of the two spectra, namely b_2_, b_3_, b_4_, b_7_, and y_1_ for the Orbitrap spectrum and y_2_, y_4_, and b_10_ for the timsTOF spectrum.

**Figure 1:**
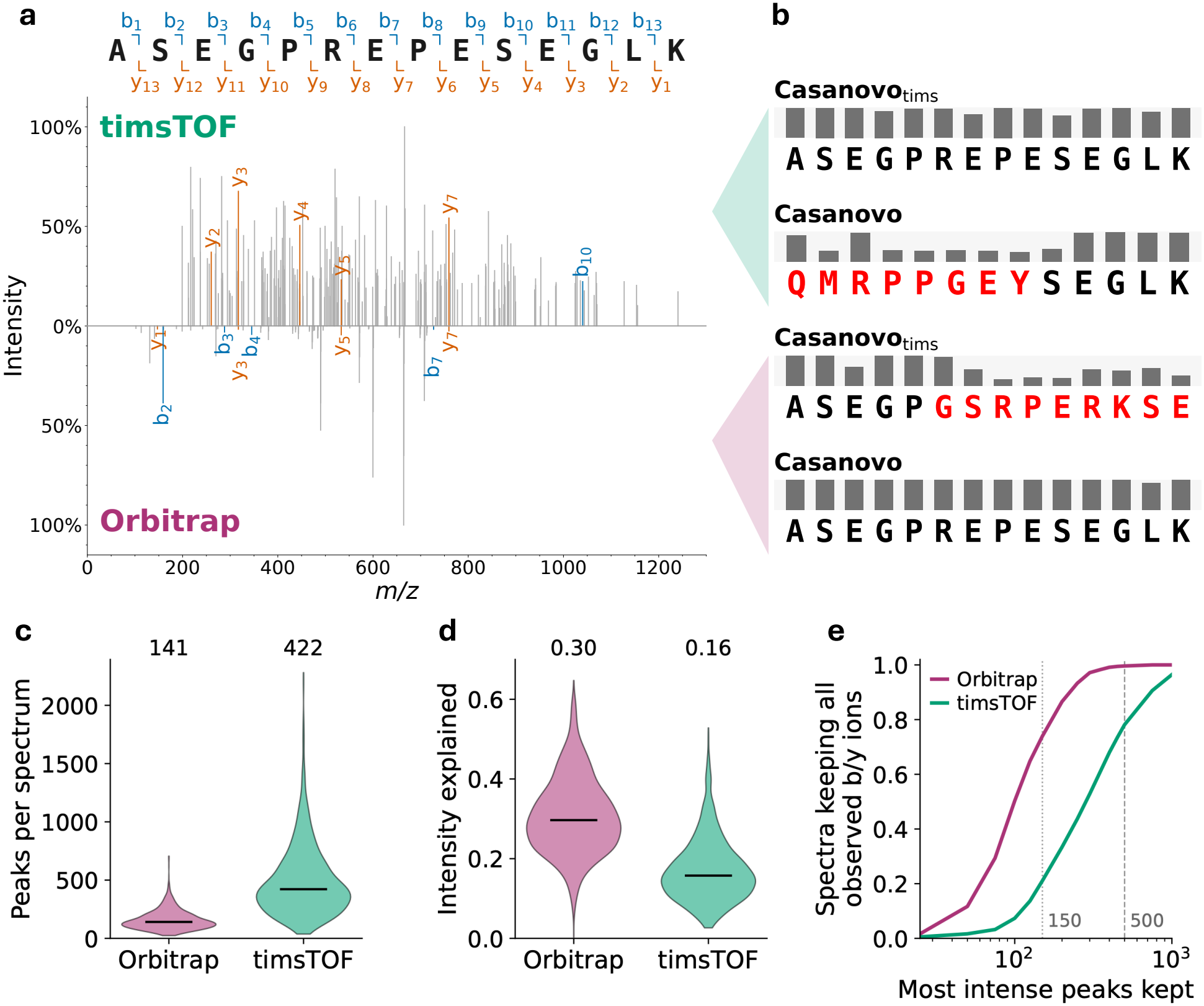
**(a)** Mirror plot of a timsTOF (top) and an Orbitrap (bottom) spectrum of the same peptide. The b and y ions of the ground-truth sequence are annotated and colored, peaks without an annotation are grey; the precursor peak and its isotopes are removed and peaks below 1% of the base peak are dropped. **(b)** Casanovo_tims_ and Casanovo predictions for both spectra of panel (a), with the per-residue confidence scores shown above each sequence. Black and red letters mark residues that are mass-correct and incorrect with respect to the ground truth, respectively. **(c)** timsTOF spectra contain about three times as many peaks as Orbitrap spectra of the same peptide (median 422 against 141). **(d)** A smaller share of that intensity is explained by the b and y ions of the detected peptide (0.16 against 0.30). **(e)** Keeping only the most intense peaks therefore costs fragment evidence far more often on timsTOF: the 150-peak limit Casanovo applies by default keeps all observed b and y ions in 21% of the timsTOF spectra, against 74% of the Orbitrap spectra, and the 500-peak limit of Casanovo_tims_ in 78% versus 99.6%. Panels (c–e) summarise 1,267 precursors from 955 peptides measured on both instruments, with one median value per combination and instrument.

Both observations hold in a systematic analysis and not just this example alone, as we compared differences on 1267 precursors from 955 peptides that were measured on both instrument types. Over all precursors, a timsTOF spectrum carries a median of 422 peaks against 141 on the Orbitrap, and the tims-TOF spectrum contains more peaks in 95.5% of the pairs (Fig. 1c). The additional peaks are, however, almost entirely unexplained signal: the share of the spectrum’s intensity that the b and y ions of the detected peptide account for drops from 0.30 to 0.16, and is lower on timsTOF in 93.5% of the pairs (Fig. 1d). The fragment ladders themselves are equally complete on both instruments, covering 0.61 and 0.58 of the possible b and y ions (Supplementary Fig. 1), so the timsTOF spectra do not contain fewer fragments. Instead, the fragments are simply harder to find among three times as many peaks.

The two instruments also differ in which fragments they record at all: the first ion of each series is effectively absent from the timsTOF spectra, with y_1_ observed in 0.1% of the precursors versus 93% on the Orbitrap and b_1_ in 0.1% against 16%, while from the fourth position inward the two instruments yield very similar y ion coverage (Supplementary Figs. 2 and 3). In addition to general differences in fragment intensity patterns, an Orbitrap-trained model therefore expects terminal-ion evidence that timsTOF spectra do not provide.

The difference in peak counts and how signal is distributed across intensity has a direct consequence for how a model must read a spectrum. Casanovo retains the 150 most intense peaks of a spectrum, which keeps all observed b and y ions in 74% of the Orbitrap spectra but in only 21% of the timsTOF spectra (Fig. 1e). Retaining 500 peaks raises the latter to 78%. The informative fragments on timsTOF therefore reach deeper into the intensity ranking than an Orbitrap-derived preprocessing setting anticipates.

Beyond preprocessing, we expect that a model trained on data from one type of instrument will not generalize well to data from a different type of instrument. While we provide a more thorough evaluation of Casanovo_tims_ in the following sections, we applied it and the original Casanovo model (version 5.2.0) to the timsTOF spectrum and the Orbitrap spectrum of Fig. 1a. The predictions (Fig. 1b) agree with our expectation: both Casanovo variants correctly sequence the peptide that was measured on their respective instrument, and both models fail to correctly predict the peptide of the opposite modality. Worse, the respective “non-native” model predicts some incorrect residues with high confidence, further diminishing applicability of the original Casanovo model in the timsTOF domain.

### Casanovo_tims_ is built from a large and diverse timsTOF dataset

Having established that timsTOF and Orbitrap spectra differ substantially in ways that affect *de novo* peptide sequencing, we next assembled a large and diverse timsTOF dataset to train Casanovo_tims_. Comparable in size to the dataset used to train the latest Casanovo model [9], it comprises over 6.7 million PSMs, 2.9 million distinct precursors, and 2.2 million unique peptides from 3,030 runs (Fig. 2a, details in Training, validation, and test data). Peptide diversity was achieved through two means. First, the runs were derived from 14 different species (Fig. 2b). Second, the runs were generated using five different enzymes, namely, trypsin, LysC, GluC, LysN, and AspN. Additionally, about one quarter of the runs come from immunopeptidomics experiments, both human leukocyte antigen (HLA) class I and class II, which exhibit non-specific cleavage. In the final dataset, approximately one-third of the runs are from non-tryptic sources, and almost 30% of the peptides are non-tryptic, i.e., they do not end in K or R (Fig. 2c). For modifications, we included methionine oxidation, cysteine carbamidomethylation, and N-terminal acetylation and ammonia loss. Additionally, besides carbamidomethylated cysteine, Casanovo_tims_ also supports unmodified cysteine and cysteinylated cysteine, which are both particularly relevant in immunopeptidomics experiments. A complete overview of the training dataset sources is provided in Table 1, and an overview of the final dataset and residue and modification abundances can be found in Supplementary Tables 1–3. The curated corpus, including the train, validation, and test splits used here, is deposited together with the scripts that generate it (Data availability), so that it can be reused as a training and benchmarking resource for timsTOF data.

**Figure 2:**
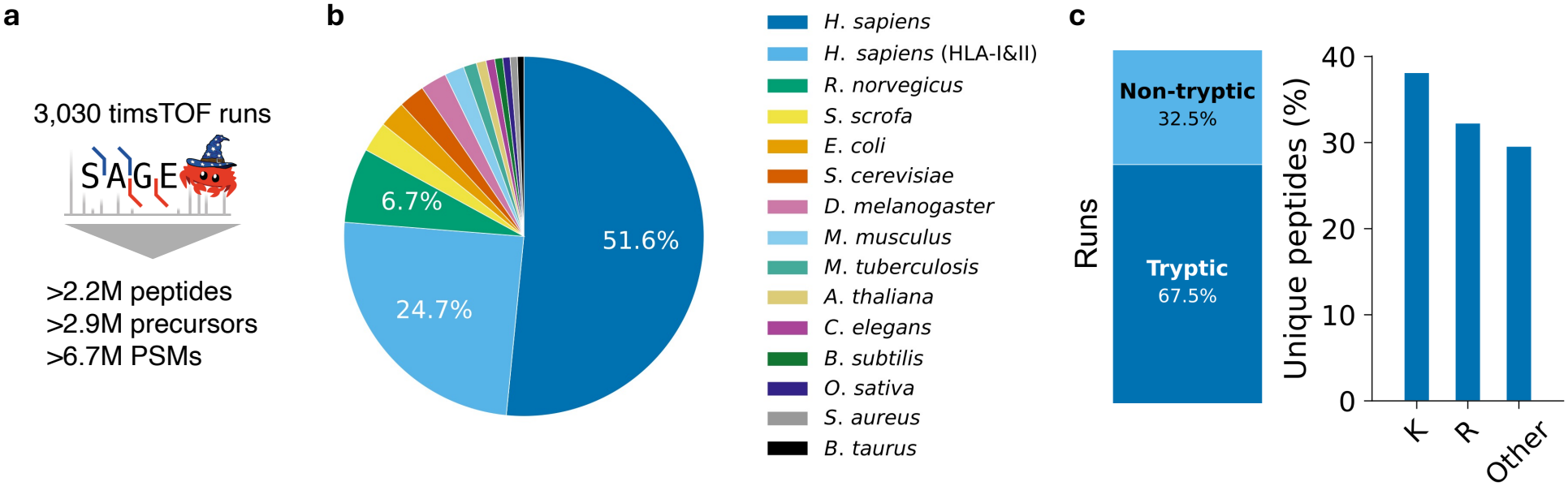
**(a)** Casanovo_tims_ is trained on a large, diverse dataset stemming from thousands of timsTOF runs that were annotated with Sage. **(b)** Data from 14 species across 3030 runs were acquired to ensure peptide diversity. **(c)** To allow Casanovo_tims_ to handle both tryptic and non-tryptic data, approximately one-third of the collected runs come from non-tryptic experiments (left). This is reflected in the peptide C-terminus statistics, where sequences not ending in K or R account for almost 30% of the unique peptides (right).

**Table 1:** Summary of datasets used for training the model. Per PRIDE project: species, protease, number of raw files, and number of annotated spectra written to either the train, validation, or test set. PT = ProteomeTools. PXD025671 was searched against a combined human + *M. tuberculosis* database; its human and *M. tuberculosis* rows therefore share the same raw files and add to the total only once.

| PRIDE project | Species | Enzyme | Raw files | Spectra |
|---|---|---|---|---|
| PXD048346 | *Arabidopsis thaliana* | Trypsin | 21 | 54,034 |
| PXD050814 | *A. thaliana* | Trypsin | 6 | 160,547 |
| PXD053191 | *Bacillus subtilis* | Trypsin | 8 | 50,188 |
| PXD053195 | *B. subtilis* | Trypsin | 4 | 41,173 |
| PXD062015 | *B. subtilis* | Trypsin | 10 | 103,673 |
| PXD030994 | *Bos taurus* | Trypsin | 18 | 22,242 |
| PXD019086 | *Caenorhabditis elegans* | Trypsin | 24 | 103,553 |
| PXD019086 | *Drosophila melanogaster* | LysC | 24 | 96,008 |
| PXD019086 | *D. melanogaster* | LysN | 24 | 144,567 |
| PXD019086 | *D. melanogaster* | Trypsin | 24 | 216,351 |
| PXD019086 | *Escherichia coli* | LysC | 25 | 18,294 |
| PXD019086 | *E. coli* | LysN | 24 | 22,788 |
| PXD019086 | *E. coli* | Trypsin | 24 | 19,797 |
| PXD019086 | *Homo sapiens* | LysC | 24 | 37,361 |
| PXD019086 | *H. sapiens* | LysN | 24 | 120,878 |
| PXD019086 | *H. sapiens* | Trypsin | 24 | 354,100 |
| PXD025671 | *H. sapiens* | AspN | 8 | 12,881 |
| PXD025671 | *H. sapiens* | GluC | 8 | 27,577 |
| PXD025671 | *H. sapiens* | Trypsin | 26 | 42,492 |
| PXD038782 (HLA1) | *H. sapiens* | None | 314 | 655,463 |
| PXD038782 (HLA2) | *H. sapiens* | None | 353 | 682,582 |
| PXD040740 (HLA1) | *H. sapiens* | None | 63 | 223,803 |
| PXD040740 (HLA2) | *H. sapiens* | None | 27 | 117,062 |
| PXD077178 | *H. sapiens* | Trypsin | 1070 | 1,207,805 |
| PXD019086 (PT/MissingGenes) | *H. sapiens* (synthetic) | Trypsin | 150 | 459,256 |
| PXD019086 (PT/Proteotypic) | *H. sapiens* (synthetic) | Trypsin | 151 | 402,739 |
| PXD019086 (PT/SRMatlas) | *H. sapiens* (synthetic) | Trypsin | 96 | 473,250 |
| PXD047626 | *Mus musculus* | Trypsin | 54 | 178,701 |
| PXD025671 | *Mycobacterium tuberculosis* | AspN | 6 | 20 |
| PXD025671 | *M. tuberculosis* | GluC | 6 | 17 |
| PXD025671 | *M. tuberculosis* | Trypsin | 24 | 300 |
| PXD029260 | *Oryza sativa* | Trypsin | 8 | 36,124 |
| PXD063170 | *O. sativa* | Trypsin | 12 | 116 |
| PXD029649 | *Rattus norvegicus* | Trypsin | 16 | 54,795 |
| PXD040661 | *R. norvegicus* | Trypsin | 62 | 15,992 |
| PXD040890 | *R. norvegicus* | Trypsin | 32 | 55,922 |
| PXD045602 | *R. norvegicus* | Trypsin | 27 | 234 |
| PXD046117 | *R. norvegicus* | Trypsin | 6 | 73,700 |
| PXD050504 | *R. norvegicus* | Trypsin | 48 | 9299 |
| PXD061633 | *R. norvegicus* | Trypsin | 15 | 23,036 |
| PXD019086 | *Saccharomyces cerevisiae* | LysC | 24 | 150,265 |
| PXD019086 | *S. cerevisiae* | LysN | 24 | 129,711 |
| PXD019086 | *S. cerevisiae* | Trypsin | 24 | 136,704 |
| PXD050156 | *Staphylococcus aureus* | Trypsin | 20 | 1496 |
| PXD057988 | *Sus scrofa* | GluC | 18 | 19,764 |
| PXD057988 | *S. scrofa* | Trypsin | 66 | 35,467 |
| **Total** |  |  | 3030 | 6,792,127 |

### Casanovo_tims_ accurately sequences peptides from timsTOF-generated spectra

After training Casanovo_tims_ on the aforementioned dataset, we evaluated it on a held-out test set of ∼680,000 timsTOF spectra derived from both tryptic and non-tryptic experiments. Critically, none of the peptides represented in the test set appeared in the training set (Training, validation, and test data). As baselines, we also applied four other models to the same test set: Casanovo, as well as InstaNovo, InstaNovo+ [2], and *π*-PrimeNovo [22] (General performance evaluation). InstaNovo is an autoregressive transformer model like Casanovo, and InstaNovo+ adds a diffusion model that iteratively refines its predictions. *π*-PrimeNovo also employs a transformer architecture but decodes non-autoregressively, emitting all residues at once under a precursor-mass constraint. All four competing methods were trained solely from Orbitrap data. To ensure a fair comparison we eliminated from the test set all spectra containing modifications that are not handled by all five models. We evaluate the performance of the models by comparing precision–coverage curves at both the peptide and amino acid levels (General performance evaluation).

Not surprisingly, training on timsTOF data allows the model to substantially outperform models that are trained solely from Orbitrap data. The difference is most pronounced at the stringent precision levels that practical application demands. At a peptide-level precision of 90%, Casanovo_tims_ provides predictions for 54.3% of the test spectra, four times the coverage of the best Orbitrap-trained baseline (*π*-PrimeNovo 13.3%, InstaNovo 12.9%, Casanovo 8.1%, and InstaNovo+ 7.6%; Fig. 3a). At 95% precision it still covers 44.0% of the spectra, a regime the baselines barely reach: InstaNovo covers 5.2%, *π*-PrimeNovo 3.3%, InstaNovo+ 1.9%, and Casanovo none at all. Conversely, when no threshold is applied and every spectrum has to be sequenced, Casanovo_tims_ is correct for 55.8% of them, against 30.7% for *π*-PrimeNovo, 28.6% for InstaNovo+, 27.3% for InstaNovo, and 18.8% for Casanovo.

**Figure 3:**
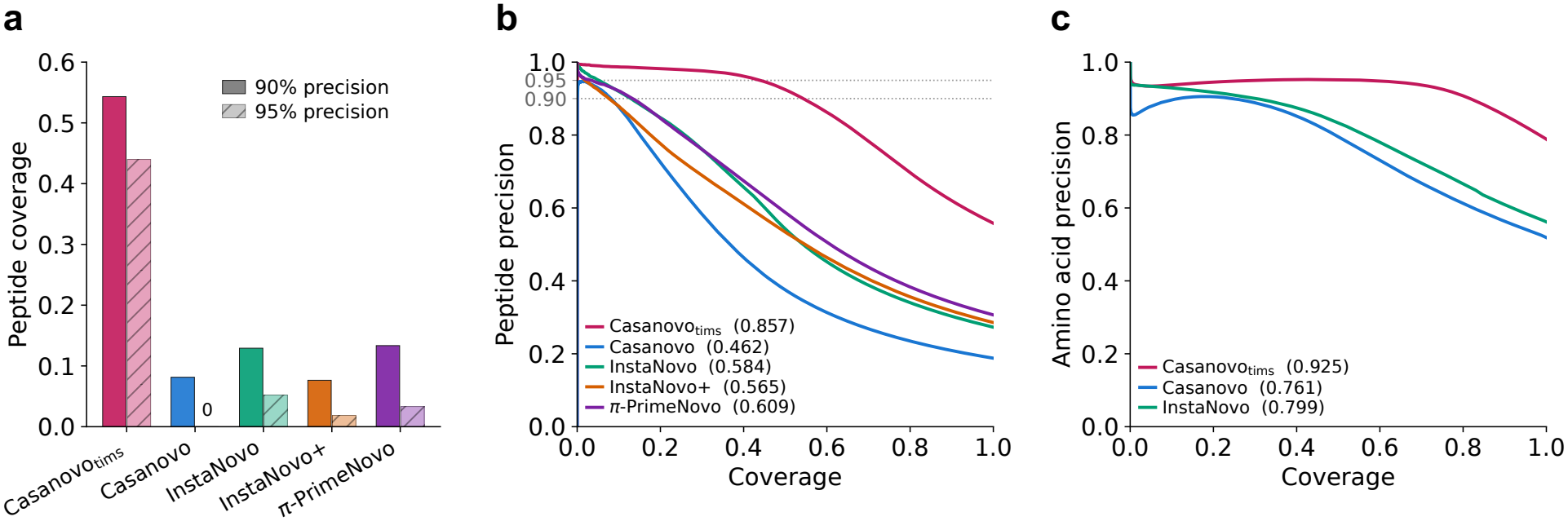
**(a)** Share of the test spectra that each model can sequence while holding a peptide-level precision of 90% and of 95%. Casanovo does not reach 95% precision at any score threshold, hence the zero. **(b)** Peptide-level precision–coverage curves on the timsTOF test set, with the average precision of each curve given in parentheses. The dotted lines mark the two precision levels of panel (a). **(c)** The same at the amino acid level. InstaNovo+ and *π*-PrimeNovo are absent because they do not report amino acid scores.

The same ordering holds over the whole operating range. At the peptide level (Fig. 3b), Casanovo_tims_ achieves an average precision (AP) of 0.857, which is substantially larger than the AP achieved by Casanovo (AP=0.462), the two variants of InstaNovo (AP=0.584 and AP=0.565, respectively), and *π*-PrimeNovo (AP=0.609). In this setting, adding the diffusion post-processor to InstaNovo appears to moderately decrease its performance. Importantly, running Casanovo on 500 peaks (instead of the default 150), which in principle includes more fragment evidence (Fig. 1e), does not improve its performance (AP=0.450, not shown). This indicates that the performance gap is due to a general domain shift between Orbitrap and timsTOF data. At the amino acid level we observe similar trends, with Casanovo_tims_ achieving an AP of 0.925 versus the next best performer (InstaNovo) at AP=0.799, and predicting 81.7% of all residues at 90% precision, compared with 30.3% for InstaNovo and 24.6% for Casanovo (Fig. 3c). InstaNovo+ and *π*-PrimeNovo are not included in this comparison because they do not report scores at the amino acid level.

Because downstream analyses often apply a score threshold rather than using the full output, we also asked what such a threshold delivers. Applying a cutoff of 0.7 retains 43.0% of the spectra at a peptide-level precision of 0.953, and the residues of those predictions are 98.3% correct (Supplementary Fig. 4); the score is informative but under-confident, so it should not be interpreted as a probability of being correct. The predictions that remain incorrect at that threshold are usually related sequences. More than half differ from the true peptide by exactly two residues, the signature of a swap or of a single isobaric substitution (Supplementary Fig. 5), and the great majority preserve the measured mass: 87% have the same mass as the precursor, 73% the same length as the true peptide, and 36% use exactly the same residues in a different order (Supplementary Fig. 6). The errors are also not spread evenly along the peptide. Consistent with the missing terminal fragments, the N-terminal residue is the least reliable position (2.5% error against 1.5% in the interior), while the C-terminal residue is the most reliable (0.7%), where the enzymatic specificity of the C terminus constrains the prediction even though y_1_ is almost never observed (Supplementary Fig. 7).

Overall, these results demonstrate that Casanovo_tims_ is able to accurately sequence unseen peptides from spectra generated by a timsTOF instrument, and that training on timsTOF data substantially improves performance relative to existing models trained solely on Orbitrap data.

### Casanovo_tims_ finds plausible neoantigen candidates in immunopeptidomics

Immunopeptidomics is a promising application area for *de novo* peptide sequencing, because it can facilitate the discovery of neoantigen candidates that are not present in reference databases. In this setting, timsTOF instruments play an important role because they are tailored toward low-input samples, which are common in immunopeptidomics. To assess the performance of Casanovo_tims_, we applied it to sequence the immunopeptidome from an HLA class I immunoprecipitated A375 melanoma cell line run, which was not seen during training (see Immunopeptidomics data).

We compared the peptides predicted by Casanovo_tims_ at high confidence (score *>*0.7, a threshold at which the model reaches a peptide-level precision of 0.953 on held-out data; Supplementary Fig. 4) to those identified by Sage at a 1% peptide-level false discovery rate (FDR) threshold and found substantial overlap (Fig. 4a). In addition to 2326 peptides detected by both methods, Casanovo_tims_ predicts 205 peptides from the canonical human proteome and 280 peptides that are non-canonical (*de novo*) and thus potential neoantigen candidates. Overall, combining Casanovo_tims_ with Sage results increases the number of unique peptides found by 15.4%.

**Figure 4:**
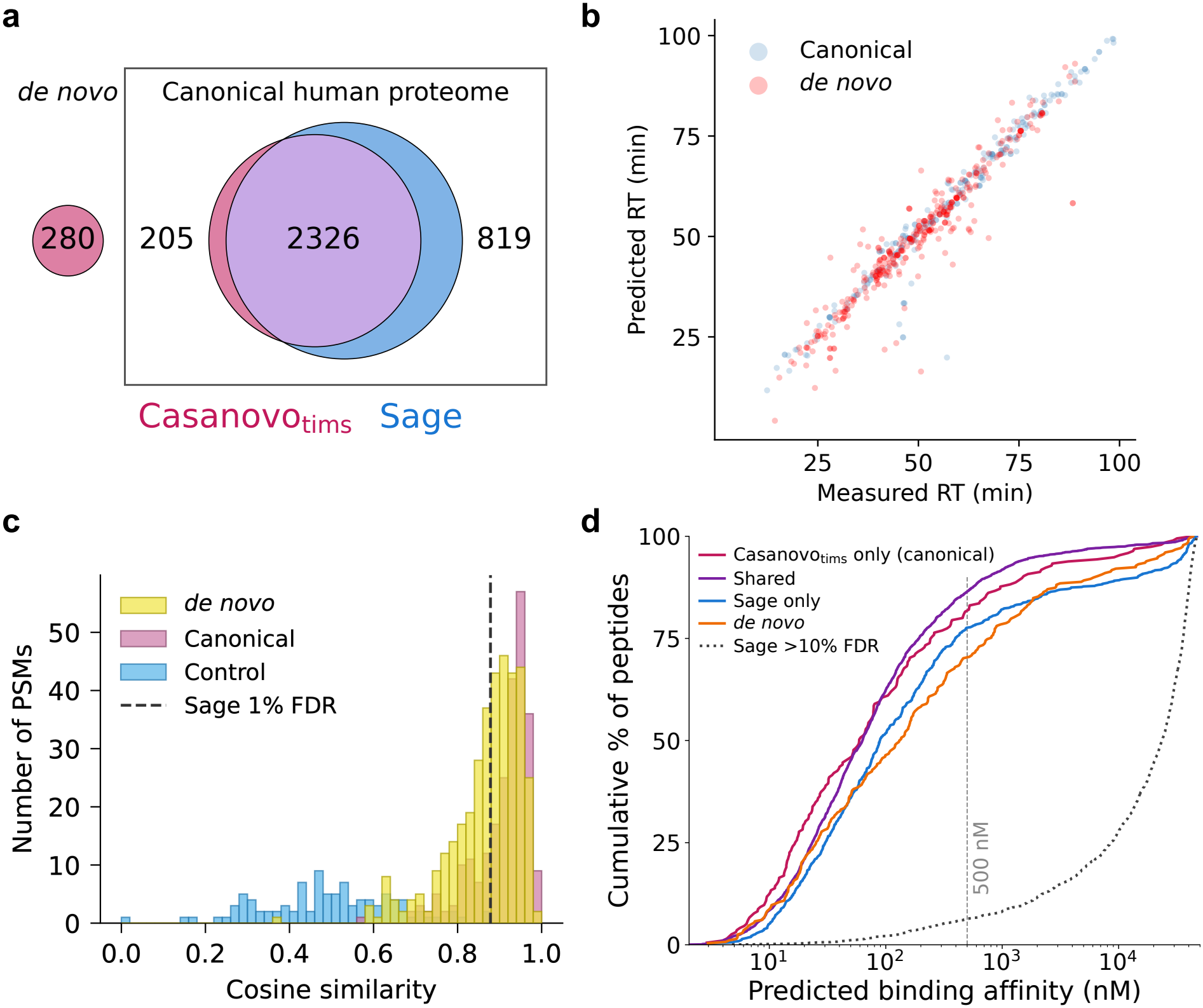
**(a)** Venn diagram indicating unique peptides found by Casanovo_tims_ and Sage in the immunopeptidomics run, including non-canonical peptides found by Casanovo_tims_. **(b)** Comparing predicted RT with observed RT for peptides found uniquely by Casanovo_tims_. **(c)** Distributions of spectral similarity between predicted and observed spectra for peptides found uniquely by Casanovo_tims_, compared to Sage predictions at 1% FDR and a control set of low-confidence Sage predictions (*>*10% FDR; a random subsample of 100 of the 1000 control PSMs is shown). **(d)** Predicted binding affinity against share of peptides found for the peptide sets introduced in (a), together with the low-confidence Sage control.

To evaluate the plausibility of the *de novo* peptides, we applied three different types of peptide property comparisons (Immunopeptidomics evaluation). First, we compared the RT predicted by AutoRT with the observed RT for each PSM from both sets, canonical and *de novo* (Fig. 4b). We observe a strong correlation for both types of peptides (Pearson *r* = 0.977 for canonical and *r* = 0.949 for *de novo*). In contrast, a control set of 1000 low-confidence Sage matches (*>*10% FDR, not pictured) shows no correlation (*r* = −0.180), confirming that the observed correlations are specific to confident detections. Because the *de novo* sequencing algorithm does not have access to information about RT, this comparison supports the plausibility of the 280 *de novo* peptides found by Casanovo_tims_.

A second means of confirming peptides predicted by Casanovo_tims_ is to compare the predicted fragment ion intensities (using MS2PIP) with the observed ones for each PSM (Fig. 4c). Again, we investigated the human and *de novo* peptide sets defined above, and we also included a control set consisting of 1000 random PSMs from Sage with an FDR of 10% or higher. While Sage detections at 1% FDR display a median cosine similarity of 0.878 (dashed vertical line), both Casanovo_tims_ high-confidence prediction sets, canonical and *de novo*, show higher median cosine similarities of 0.933 and 0.888, respectively. Thus, Casanovo_tims_’s generated peptides are at least as consistent with the observed spectra as the high-confidence Sage predictions.

As a more direct test of whether the detected peptides are plausible HLA ligands, we predicted the binding affinity of each peptide against the sample’s six-allele HLA class I genotype using NetMHCpan-4.2, taking the strongest predicted affinity across alleles (Immunopeptidomics evaluation). As before, we considered Casanovo_tims_ predictions with a score greater than 0.7 and Sage detections at 1% FDR, and we used low-confidence Sage matches (*>*10% FDR) as a negative control. Shown in Fig. 4d, shared peptides were strongly enriched for predicted binders: 86.5% fell below the 500 nM threshold and 45.1% were strong binders (*<*50 nM), with a median affinity of 63 nM. Peptides detected only by Casanovo_tims_ showed an equally favorable profile, with 81.5% below 500 nM, 45.6% strong binders and a median affinity of 59 nM. This observation suggests that peptides recovered by Casanovo_tims_ but missed by database search largely reflect true immunopeptidome coverage. The *de novo* peptides, which do not match the reference proteome, were also mostly predicted binders (70.1% below 500 nM, median 133 nM). Sage-only peptides were somewhat weaker binders (77.6% below 500 nM, median 89 nM), while the low-confidence control looked like non-binding background, with only 6.3% below 500 nM and a median affinity of roughly 2.4 × 10^4^ nM, confirming that the binding signal is specific to confident detections.

Peptide length provides another, prediction-free check. HLA class I ligands are overwhelmingly nine residues long, and the Casanovo_tims_ predictions follow that distribution closely: 78% of the Casanovo_tims_-only peptides and 63% of the *de novo* peptides are 9-mers, against 70% of the peptides both methods agree on. The low-confidence control shows no such preference (22% 9-mers, 21% 8-mers, and 26% 10-mers), as expected for matches that are not genuine ligands.

These results demonstrate that Casanovo_tims_ is a powerful tool in immunopeptidomics, capable of detecting neoantigen candidates that are not found by database search methods.

### Casanovo_tims_ can decipher samples from complex microbial communities

In metaproteomics, samples of interest are often derived from complex microbial communities, such as the human gut microbiome, where the diversity of organisms and the presence of uncharacterized species can limit the effectiveness of database search methods. Casanovo_tims_ provides a complementary approach by enabling *de novo* peptide sequencing, which does not rely on existing protein databases.

Here, using timsTOF data from a human gut microbiome sample (Metaproteomics data), we compared Casanovo_tims_ predictions to those obtained from a Sage database search against the UHGP-90 catalogue, which contains 13.8 million protein clusters derived from the human gut microbiome. Casanovo_tims_ (using no confidence threshold) recovers 8187 unique peptides that are present in UHGP-90, compared with Sage’s 10,537. We restricted Casanovo_tims_ predictions to peptides of 10–50 residues and excluded low-complexity sequences. A lookup against a decoy catalogue matches 39 of these predictions, or 0.48% of the 8187, implying that the error rate of the catalogue matching is well below the 1% peptide-level FDR at which the Sage results are reported (Metaproteomics evaluation). Importantly, the two methods agree on the majority of peptides, with 6539 shared between both methods and Casanovo_tims_ finding 1648 additional peptides not found by Sage (Fig. 5a).

**Figure 5:**
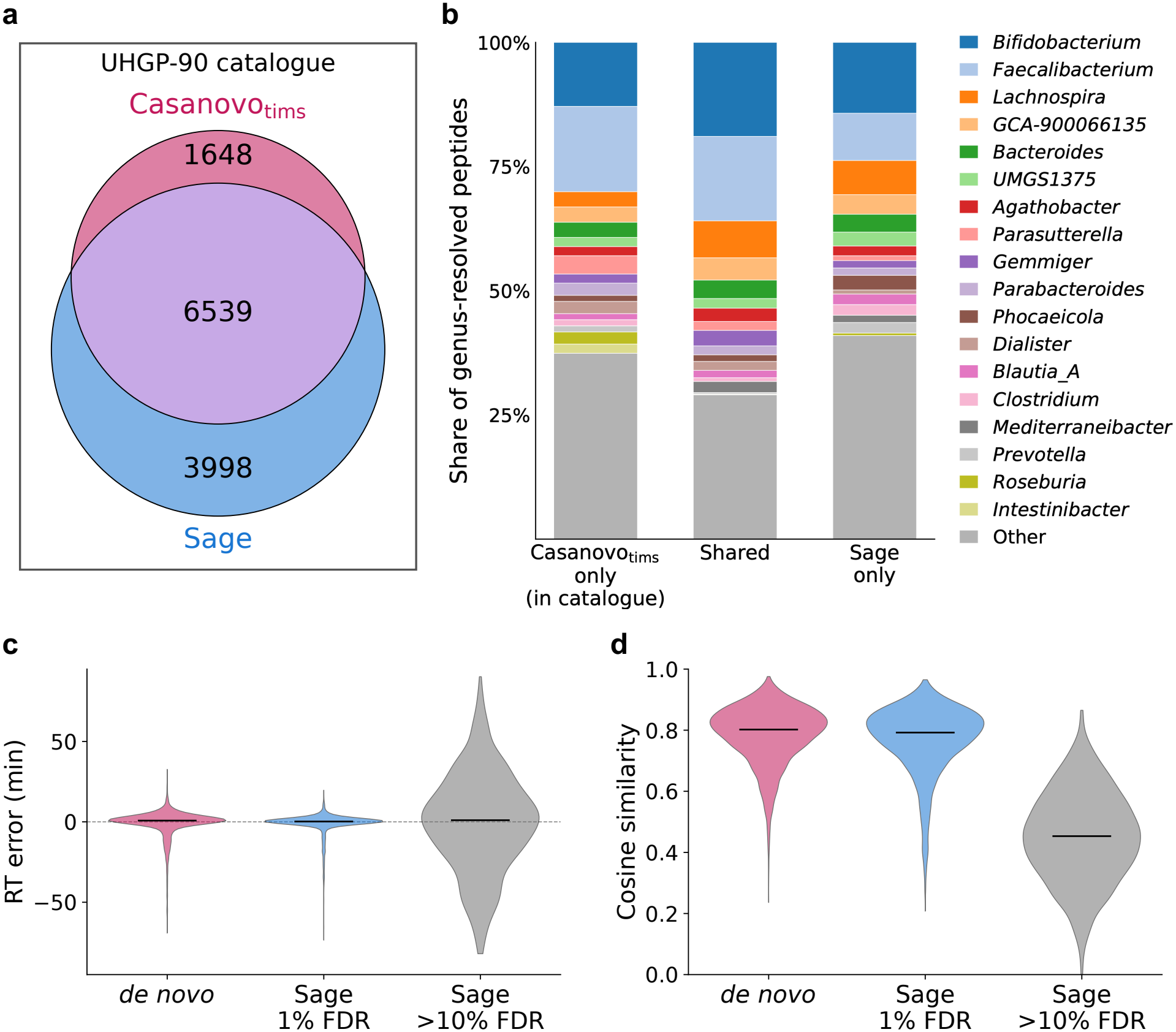
**(a)** Venn diagram indicating the number of unique peptides detected by Casanovo_tims_ and Sage in the UHGP-90 catalogue. **(b)** Genus distribution of peptides in UHGP-90 detected uniquely by Casanovo_tims_ (left), both Casanovo_tims_ and Sage (center), and Sage only (right). **(c)** Violin plot comparing predicted RTs (using AutoRT) with observed RTs for *de novo* peptides predicted by Casanovo_tims_, Sage detections at 1% FDR, and a low-confidence Sage *>*10% FDR control set. **(d)** Violin plot comparing cosine similarities between predicted (using MS2PIP) and observed spectra for *de novo* peptides predicted by Casanovo_tims_, Sage detections at 1% FDR, and a low-confidence Sage *>*10% FDR control set.

We explore these three sets—Casanovo_tims_-only, shared, and Sage-only—further in Fig. 5b, where we show the genus-level taxonomic composition of the peptides detected by each method (Metaproteomics evaluation). All three distributions generally agree on the most abundant genera (i.e., *Bifidobacterium, Faecalibacterium, Lachnospira*). These results suggest two conclusions: (i) Casanovo_tims_ is able to capture a broad representation of the microbial community without relying on a reference database, and (ii) when using database search, Casanovo_tims_ can detect additional peptides that are not found by Sage and thus provide further evidence of the microbial community composition.

Additionally, we conducted an analysis of *de novo* peptides predicted by Casanovo_tims_ that are not found in the UHGP-90 catalogue. Similar to the immunopeptidomics analysis, we compared the predicted RT and fragment ion intensities to the observed values for these peptides, using a Casanovo_tims_ confidence threshold of 0.7 (Metaproteomics evaluation). Overall, Casanovo_tims_ generates 2688 unique peptides from 4388 PSMs that are not in UHGP-90. We compare these to 2397 unique peptides from 4210 PSMs from the 1% FDR Sage predictions that were not used for finetuning AutoRT, and to 929 PSMs that the Sage search rejected (*>*10% FDR) as a negative control. The RT errors (predicted minus measured, Fig. 5c) are centered on zero for both sets, with a slightly wider spread for the *de novo* set (median absolute error 2.32 vs. 1.19 min; Pearson *r* = 0.9447 vs. 0.9654). The control is centered on zero as well but scatters far more widely (18.92 min; *r* = 0.3679). Predicted and measured spectra agree well for both sets (Fig. 5d; median cosine similarity 0.802 vs. 0.792), with a slight advantage for the *de novo* set, and clearly better than the control (0.453; 37.1% of its PSMs above 0.5, against 98.8% and 96.1%).

Overall, these results suggest that Casanovo_tims_ is able to generate plausible peptide predictions that are not found in existing catalogues and thus may represent novel peptides from uncharacterized species in metaproteomic samples.

## Discussion

Because timsTOF instruments are becoming more widely used in proteomics, we used the increasing availability of timsTOF data to curate a large-scale timsTOF dataset to train Casanovo_tims_, a transformer-based model for *de novo* peptide sequencing. The resulting model strongly outperforms existing models trained on Orbitrap data, achieving higher AP at both the peptide and amino acid levels. At 95% peptide-level precision, Casanovo_tims_ reaches an eightfold increase compared with the strongest Orbitrap-trained baseline. Additionally, we showed that Casanovo_tims_ is able to detect plausible neoantigen candidates in immunopeptidomics data and provides complementary peptide detections in metaproteomics data.

Casanovo_tims_ still has limitations: it does not support phosphorylation or other, potentially rare post-translational modifications. This can be attributed to the fact that, compared to Orbitrap data, timsTOF data is still relatively scarce, especially in terms of datasets enriched for modifications. Here, a transfer learning approach could be beneficial, where a model is first trained on a wide range of modifications from Orbitrap data and then finetuned on a smaller set of timsTOF data with the same modifications.

In addition, Casanovo_tims_ was only trained on ddaPASEF data. Because data-independent acquisition (DIA) methods generally produce more complex, chimeric spectra compared to data-dependent acquisition methods, we expect Casanovo_tims_ applicability to this type of data to be limited. A model similar to Cascadia [23] but trained on diaPASEF data could be beneficial, as it would be able to learn the specific characteristics of DIA spectra and improve peptide sequencing performance.

A general limitation of *de novo* peptide sequencing is that stringent FDR control for *de novo* peptides is missing. While the presented peptide property analyses suggest that the majority of high-confidence *de novo* peptides are plausible, we cannot provide a rigorous FDR estimate for them like database search methods can. Introducing such an FDR control mechanism would improve applicability of *de novo* peptide sequencing tools such as Casanovo_tims_ and is an important direction for future work.

The peptides that Casanovo_tims_ recovers beyond the database search are supported by several independent lines of evidence, but all of them are predictions rather than measurements. Synthesizing a subset of the *de novo* peptides and re-acquiring them would convert this plausibility argument into direct confirmation, and is the natural next step for the neoantigen candidates in particular.

Overall, Casanovo_tims_ closes the gap between Orbitrap-trained *de novo* models and timsTOF data, making *de novo* peptide sequencing a practical option on an instrument class where it previously was not. Given the model’s accuracy on timsTOF data, it can be used as a complement to database search in applications where reference databases are incomplete.

## Methods

### Casanovo_tims_ model

Casanovo is a deep learning model that translates from an observed MS/MS spectrum to a corresponding amino acid sequence. The model employs a transformer architecture [24] with a standard encoder–decoder structure. The encoder takes as input a sequence of peaks, represented as (*m/z*, intensity) tuples. The decoder operates in an autoregressive fashion, meaning that the peptide sequence is generated one amino acid at a time. Casanovo_tims_ uses the existing Casanovo hyperparameter configuration with two changes. First, we use the top 500 peaks instead of top 150 (in terms of intensity) to account for the more complex timsTOF spectra. We compared different settings (top 150, 300, 500, and 750 peaks, Supplementary Fig. 8) and found that the top 500 peaks provide a good trade-off between accuracy and inference time (Supplementary Fig. 9). Second, we modify the vocabulary of allowed post-translational modifications, adding cysteinylation, allowing cysteine to occur unmodified, and excluding four variable modifications (deamidation of asparagine, deamidation of glutamine, peptide N-terminal carbamylation, and peptide N-terminal carbamylation with ammonia loss). Additionally, we have conducted an analysis that included ion mobility as an additional precursor feature, but found that this feature did not improve performance (Supplementary Fig. 10). Native timsTOF support was added to depthcharge (https://github.com/wfondrie/depthcharge), which Casanovo builds upon. More details about the Casanovo model and implementation are provided in the original publications [7, 8] and GitHub repository (https://github.com/Noble-Lab/casanovo). Casanovo_tims_ was trained for 48 epochs on a single NVIDIA H100 GPU with a batch size of 392.

### Data

#### Training, validation, and test data

To train and validate our model, timsTOF data were aggregated from 3030 runs produced using five different proteases, plus non-enzymatic immunopeptidomics runs. The data are derived from 14 species and 23 PRIDE projects (Table 1). The respective proteome databases were downloaded from UniProt [25] (https://www.uniprot.org). The data were processed in three stages: searching and rescoring, deamidation removal, and redundancy removal.

Each run was searched against its respective reference proteome with Sage [26], allowing one static modification, carbamidomethylation of cysteine (+57.0215 Da), and seven variable modifications: methionine oxidation (+15.9949 Da), asparagine and glutamine deamidation (+0.9840 Da), and peptide N-terminal acetylation (+42.0106 Da), carbamylation (+43.0058 Da), ammonia loss (−17.0265 Da), and combined carbamylation with ammonia loss (+25.9793 Da). Peptide lengths were restricted to 7–50 residues. Typically, Sage carries out PSM rescoring internally; however, when necessary due to computational restrictions, some runs were split into chunks, each searched by Sage, and then rescored by mokapot (v0.10.0) [27]. We used Sage version 0.15.0-beta.1, a pre-release build provided by Bruker that applies the vendor’s official timsTOF calibration. This build was not publicly available at the time of writing; the public Sage implements the same search algorithm, and we therefore expect comparable results, although we did not systematically compare the two.

Data from immunopeptidomics runs were searched somewhat differently. For HLA class I runs, carbamidomethylation was moved from the static to the variable modifications and cysteinylation of cysteine (+119.0041 Da) was added, with no static modifications. For HLA class II immunopeptidomics runs, only cysteinylation and carbamidomethylation of cysteine, methionine oxidation, and N-terminal acetylation were used as variable modifications, and no static modifications were used. Enzyme specificity was set to non-specific, and peptide lengths were restricted to 8–16 residues for HLA class I, and were restricted to two length windows, 8–16 and 17–23 residues, for HLA class II. For both HLA class I and II runs, the fasta files were split into smaller batches, searched separately and in the end rescored by mokapot. After the search, PSMs with carbamylation or combined carbamylation with ammonia loss were discarded due to their low abundance.

After searching, deamidations were validated and then removed. For each deamidated PSM, we applied a 5 ppm mass filter to remove misassigned isotope errors [28]. Subsequently, deamidated asparagine and glutamine were replaced by aspartic acid and glutamic acid, respectively.

Finally, PSM redundancy was reduced in several steps. First, per unique precursor (i.e., peptide and associated charge state) we ranked matches by q-value, and we downsampled the data by selecting up to three PSMs per precursor. Only PSMs with a q-value of less than 0.01 were considered. Where possible, these PSMs were required to come from different experiments; failing that, from different runs within an experiment; and failing that, from different spectra within a run. To decrease spectrum complexity, the ion mobility dimension was collapsed during spectrum extraction, yielding one fragment spectrum per precursor, which was then matched to the Sage results.

The resulting dataset consists of 6,792,127 PSMs, 2,981,749 distinct precursors, and 2,292,408 unique peptides. These data were split for training, validation, and testing in a ratio of 8:1:1 at the peptide level. The validation set was used to monitor training progress, prevent overfitting, and to select the best model checkpoint. For evaluation, we removed from the test set every PSM whose peptide (including modifications), after replacing isoleucine with leucine, also occurs in the training set. The configuration files used for the Sage searches are available as Supplementary Data 1.

#### Immunopeptidomics data

For validation on immunopeptidomics data, we applied Casanovo_tims_ to a run unseen during training from HLA class I peptide sequencing derived from A375 melanoma cell lines [29], where HLA class I receptors and the bound peptides were lysed from the cell surface and immunoprecipitated to extract the peptides. A Bruker timsTOF SCP instrument was used with an optimized single-shot ddaPASEF technique. More details regarding instrumentation and sample preparation are provided in the original publication. For our evaluation, we used the run `E_20221201_NO30_400nL_HLAc1_5e6_directIP_titration_rep3_Slot2-2_1_3511.d`. The run was searched with Sage with the same settings as the other HLA class I runs.

#### Metaproteomics data

Casanovo_tims_ was applied to a metaproteomics run from Gao *et al*. [30] (PXD063844, iProX IPX0011920000), namely `20240423_BC101_Slot1-13_1_9755.d`. These spectra were searched with Sage against the Unified Human Gastrointestinal Protein catalogue clustered at 90% sequence identity (UHGP-90; 13.8 million protein clusters) [31, 32]. The same search settings as for training data construction were applied, with the exception of doing a semi-tryptic search to account for peptides generated by in-source fragmentation [33]. Due to the large database size, the fasta file was split, and intermediate PSMs were rescored and filtered to a 1% FDR with mokapot.

### Evaluation

#### General performance evaluation

We applied `casanovoutils` (https://github.com/Noble-Lab/casanovoutils) to evaluate the performance of Casanovo_tims_ and the baseline models on the test set. Performance is evaluated using precision–coverage curves. To create a precision–coverage curve at the PSM level, spectra are sorted by decreasing Casanovo score; for every cutoff, coverage is defined as the fraction of all spectra above the threshold and precision is defined as the fraction of those spectra that are correctly predicted. The curve plots precision as a function of coverage. A similar precision–coverage curve can be plotted at the amino acid level by sorting all amino acid predictions by scores. In either case, the performance measure is the area under this curve, which is equivalent to the AP. For evaluation at both the peptide and amino acid level, we allow for the substitution of isoleucine with leucine (I/L), lysine with glutamine (K/Q), and phenylalanine with oxidized methionine (F/M[Oxidation]). Each predicted peptide is deemed correct if it matches the ground truth peptide. For the amino acid level, if ground truth and predictions are of different lengths, then the shorter sequence is padded in a way that maximizes the number of matching amino acids. An amino acid is then correct if it matches the ground truth. As baselines, we used Casanovo (version 5.2.0), InstaNovo and InstaNovo+ (version 1.2.2), and *π*-PrimeNovo (no version specified, commit hash: fc6b0035741de98af870b0b1afced7c727711e6e).

#### Immunopeptidomics evaluation

Casanovo_tims_ predictions were evaluated by comparing predicted RT and fragment ion intensities to the observed values. Only Casanovo_tims_ predictions with a peptide length of at least 8 amino acids were considered for this evaluation. As a negative control, 1000 Sage matches with peptide- and spectrum-level q-values above 0.1 were randomly sampled. For the RT comparison, peptides detected by Sage at a 1% peptide-level FDR threshold were split 80/20; the 80% split was used to fit the RT predictor, and the held-out 20% was evaluated alongside the Casanovo_tims_ predictions. For RT, we finetuned AutoRT v1.0 [34] on the 80% split (averaged RT across replicated peptides). AutoRT was run with the following parameters: `-e 40 -b 64 -u m -rlr -n 10`. Because the spectral similarity comparison requires no fitting, all Sage detections at a 1% FDR were used as the reference. Theoretical spectra were generated using MS2PIP [18] with the timsTOF model. Predicted b- and y-ion series (*m/z* values and intensities) were concatenated into a single reference spectrum; MS2PIP intensities, which are reported in log space, were exponentiated. Both intensity vectors were first square-root transformed after scaling to their respective maxima. Peaks were then aligned by matching each predicted fragment to the experimental peak of nearest *m/z*, retaining the match only if the mass difference was within a tolerance of 0.05 Da. Unmatched experimental peaks were discarded, so the comparison vectors span the set of predicted fragments. Predicted and experimental spectra were compared using cosine similarity.

Binding affinity to HLA class I molecules was predicted with NetMHCpan-4.2 [35] in binding-affinity mode, using the six HLA class I alleles of the A375 cell line listed in Cellosaurus [36] (CVCL 0132; HLA-A*01:01, A*02:01, B*44:03, B*57:01, C*06:02 and C*16:02 [37]). Peptides were assigned to the Casanovo_tims_-only, shared, Sage-only, and *de novo* sets at the sequence level, with modifications removed and isoleucine and leucine treated as identical, and restricted to 8–14 residues. Predictions were made without flanking sequence context, because *de novo* peptides and decoy matches have no source protein. For each peptide, the strongest predicted affinity across the six alleles was used, and peptides below 500 nM and 50 nM were counted as binders and strong binders, respectively. As a negative control, 2000 peptides were randomly sampled from the Sage matches with peptide- and spectrum-level q-values above 0.1 (decoys included) that did not occur in any other set.

#### Metaproteomics evaluation

For validation using metaproteomics data, the peptide sequences predicted by Casanovo_tims_ were matched against the UHGP-90 (v2.0.2) representative sequences [31, 32] by exact substring matching (pyahocorasick 2.3.1), with isoleucine and leucine both mapped to leucine. A peptide counted as present in the catalogue if it occurred in at least one representative sequence. To avoid chance matches, peptides shorter than 10 residues were discarded, as were peptides containing a subsequence of at least five identical residues or in which a single residue made up at least 50% of the sequence. Peptides were then assigned to taxa by a lowest-common-ancestor approach. Because UHGP-90 clusters proteins at 90% sequence identity, each matched representative was expanded to all members of its cluster, and the source genome of each member was taken from its protein identifier. Each genome was annotated with its Genome Taxonomy Database [38] lineage as provided in the Unified Human Gastrointestinal Genome catalogue v2.0.2 metadata; genomes without a lineage were ignored. For each peptide, the lineages of all genomes containing it were compared rank by rank from domain to species, and the peptide was assigned to the deepest rank at which they agreed, together with the corresponding consensus lineage. Peptide property comparisons were performed in the same way as for the immunopeptidomics data (Immunopeptidomics evaluation). For the catalogue-membership criterion, we repeated the UHGP-90 lookup against a decoy catalogue for error estimation. Decoy proteins were generated by reversing the residues between cleavage sites while leaving every lysine and arginine in place. Both lookups used the same predictions, the same exact-match criterion with isoleucine and leucine treated as equivalent, and the filters as introduced above.

## Acknowledgements

This work was supported by National Science Foundation award 2245300. W.B. acknowledges support by the Research Foundation – Flanders (FWO G087625N).

## Author contributions

|  | MD | BK | SW | JS | FJT | WB | GR | WSN |
|---|---|---|---|---|---|---|---|---|
| Conceptualization |  |  |  |  |  | ● | ● | ● |
| Data curation | ● | ● |  |  |  |  |  |  |
| Formal analysis | ● | ● |  | ● |  |  |  |  |
| Funding acquisition |  |  |  |  | ● |  |  | ● |
| Investigation | ● | ● |  | ● |  |  |  |  |
| Resources |  |  |  |  | ● |  |  | ● |
| Software | ● | ● | ● |  |  | ● |  |  |
| Supervision |  |  |  |  | ● | ● | ● | ● |
| Validation | ● | ● | ● | ● |  |  |  |  |
| Visualization | ● | ● |  |  |  |  |  |  |
| Writing—original draft | ● | ● |  |  |  | ● |  | ● |
| Writing—review & editing | ● | ● |  |  | ● | ● | ● | ● |

## Data availability

All data used in this study are publicly available; accessions are listed in Table 1 and in the Methods. Data used to train and validate Casanovo_tims_, the scripts to produce them, and instructions on how to reproduce the results of this study are available at https://zenodo.org/records/23592958.

## Code availability

Casanovo source code, as well as model weights, are available with an Apache 2.0 license at https://github.com/Noble-Lab/casanovo/releases.

