## Supplementary Information for "Instrument-specific learning improves *de novo* peptide sequencing from timsTOF spectra"

### Supplementary Figures

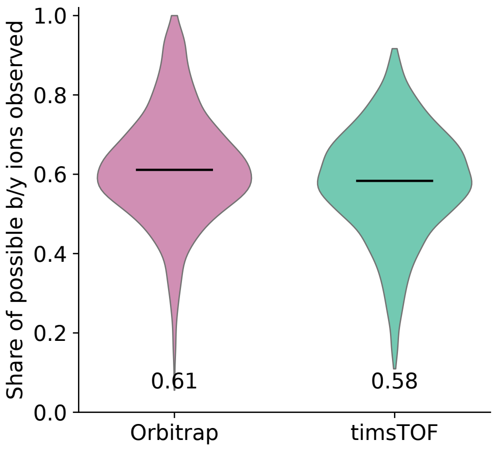


Supplementary Figure 1: **Fragment ladders are equally complete on both instruments.** The share of the $2(n-1)$ possible b and y ions of a peptide that are observed, for the 1,267 precursors, from 955 peptides measured on both instrument types, that are summarised in Fig. 1, with one median value per precursor and instrument. The medians are 0.61 on the Orbitrap and 0.58 on timsTOF, so timsTOF spectra do not contain fewer fragments of the identified peptide; they contain them among roughly three times as many peaks. Spectra were preprocessed as for the mirror plot of Fig. 1, by removing the precursor peak and its first two isotopes and discarding peaks below 1% of the base peak; no limit was placed on the number of peaks.

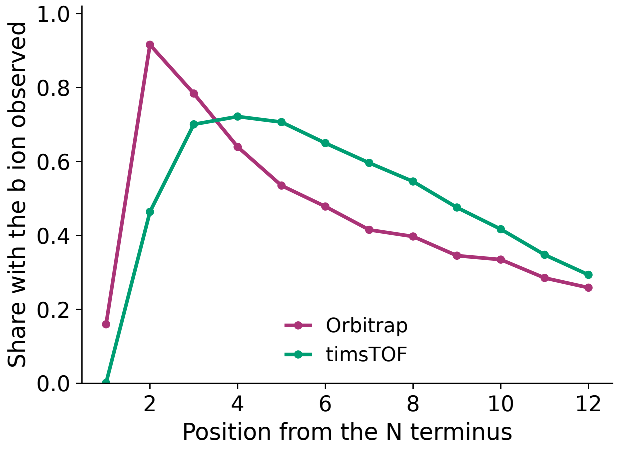


Supplementary Figure 2: **b ions by position from the N terminus.** The share of the same 1,267 precursors in which the b ion at a given position is observed. $b_1$ is effectively absent from the timsTOF spectra (0.1% against 16% on the Orbitrap) and $b_2$ is halved (46% against 92%), whereas from the fourth position inward the timsTOF spectra carry b ions more often than the Orbitrap spectra do. Spectra were preprocessed as for the mirror plot of Fig. 1, by removing the precursor peak and its first two isotopes and discarding peaks below 1% of the base peak; no limit was placed on the number of peaks.

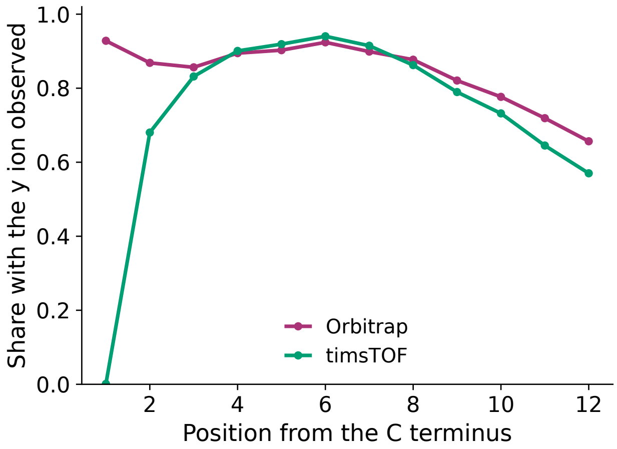


Supplementary Figure 3: **y ions by position from the C terminus.** As Supplementary Figure 2, for the y ion series. $y_1$ is observed in 93% of the precursors on the Orbitrap and in 0.1% on timsTOF, while from the fourth position inward the two instruments yield very similar y ion coverage. Fragments below $m/z$ 200 make up 1.5% of the peaks of a timsTOF spectrum against 17% of an Orbitrap spectrum, so the depletion is a property of the data as acquired. Spectra were preprocessed as for the mirror plot of Fig. 1, by removing the precursor peak and its first two isotopes and discarding peaks below 1% of the base peak; no limit was placed on the number of peaks.

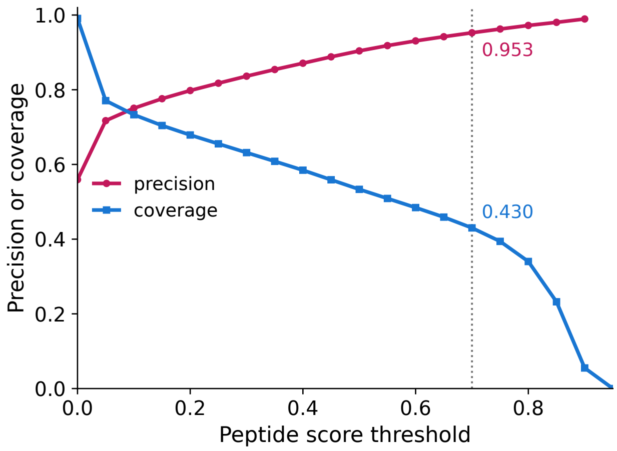


Supplementary Figure 4: **What a confidence threshold delivers.** Peptide-level precision and coverage on the held-out test set as the peptide-score threshold is varied, with the commonly used value of 0.7 marked. At that threshold Casanovo$_{\mathrm{tims}}$sequences 43.0% of the spectra at a precision of 0.953, and the residues of those predictions are 98.3% correct. Precision here is cumulative over everything kept. The score is informative but under-confident throughout, so it should not be read as a probability of being correct. Coverage is the share of all spectra in the file, so it includes the 0.9% that the model declined to sequence.

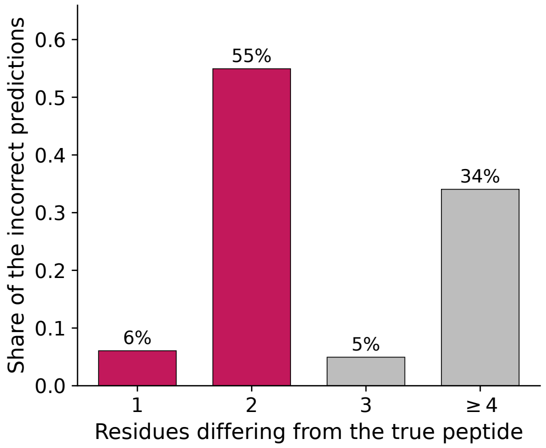


Supplementary Figure 5: **Most incorrect predictions differ from the true peptide by two residues.** The 13,667 predictions that are incorrect among the 287,901 kept at a peptide score of 0.7 (4.7%), by the number of residues that differ from the true peptide. A difference of exactly two dominates, which is the signature of a swap of two residues or of a single isobaric substitution rather than of an unrelated sequence. A third of the errors are larger than three residues, so the near misses are the rule but not the whole picture.

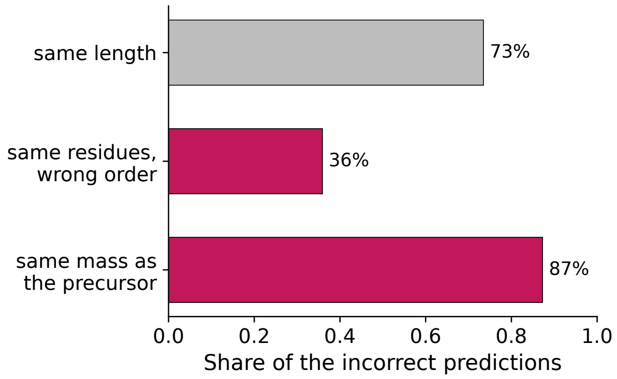


Supplementary Figure 6: **The incorrect predictions are mostly mass-preserving rearrangements.** For the same incorrect predictions as Supplementary Figure 5: the share that have the same length as the true peptide, that use exactly the same residues in a different order, and that have the same mass as the measured precursor. The model rarely proposes a peptide that the precursor mass excludes, so its errors are largely confined to the arrangement of a correct residue composition. Mass agreement is taken from Casanovo's own calculated precursor $m/z$ against the measured value, within 0.05 Da of the neutral mass.

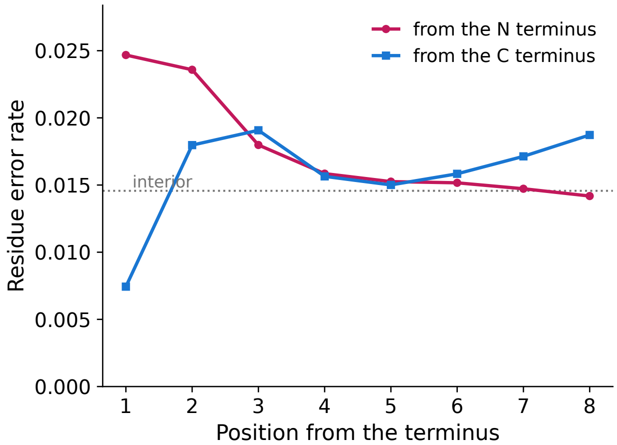


Supplementary Figure 7: **Residue errors concentrate at the N terminus.** The share of residues predicted incorrectly by position from each terminus, over the predictions kept at a peptide score of 0.7, with the rate for residues at least four positions from both ends marked. The N-terminal residue is the least reliable position at 2.5%, against 1.5% in the interior, while the C-terminal residue is the most reliable at 0.7%: the enzymatic specificity of the C terminus constrains the prediction even though $y_1$ is almost never observed (Supplementary Figure 3), whereas the N terminus has neither that constraint nor a $b_1$ ion (Supplementary Figure 2).

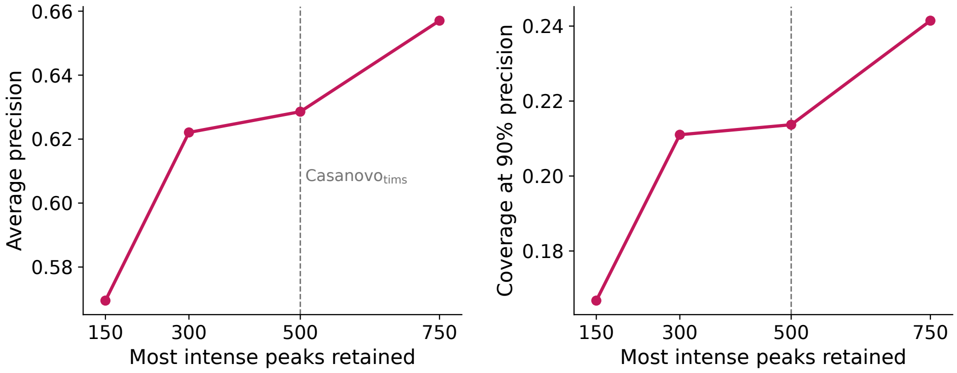


Supplementary Figure 8: **Sequencing accuracy against the number of peaks retained.** Four models trained identically except for the number of most intense peaks kept per spectrum. All four used a fixed subset of the corpus, drawn once with a fixed seed and shared by every arm: 300,000 training, 30,000 validation and 100,000 test spectra, the splits being peptidoform-disjoint so that no peptide of the test subset occurs in training. Each arm used the same architecture and schedule as Casanovo$_{\mathrm{tims}}$(512 model dimensions, 9 layers, learning rate $5 \times 10^{-4}$, weight decay $10^{-5}$, 100,000 warmup iterations, 600,000-iteration cosine period) with a batch size of 64, and was trained for 30 epochs.

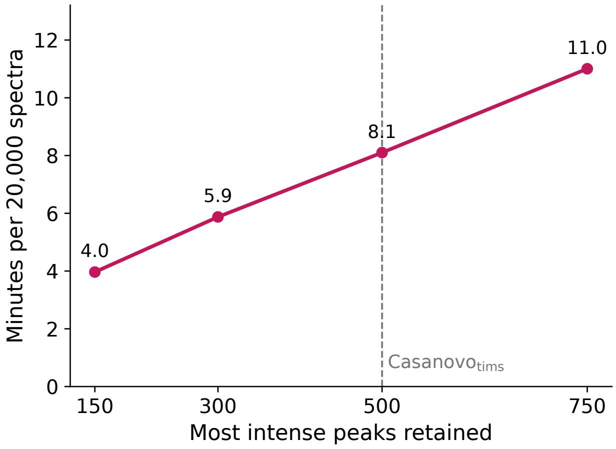


Supplementary Figure 9: **What the peak limit costs in inference time.** Time to sequence 20,000 spectra against the number of peaks retained. Raising the limit from 150 to 500 peaks, as Casanovo$_{\mathrm{tims}}$does, doubles the inference time and buys 0.059 average precision (Supplementary Figure 8); raising it further to 750 costs 1.4 times as much again for another 0.028. The four models are those of Supplementary Figure 8. All four sequenced the same 20,000 held-out spectra on the same NVIDIA A100 within a single job, after a warm-up pass that was discarded, and the times are Casanovo's own reported sequencing time, which excludes interpreter start-up and model loading. The figure compares the arms with one another; it is not a throughput specification, since it reflects one GPU, one batch size and one beam setting.

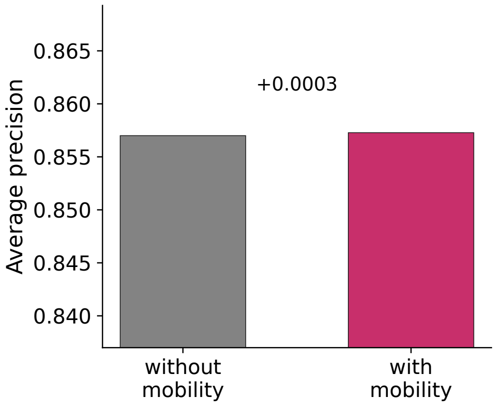


Supplementary Figure 10: **Ion mobility as an additional precursor feature.** Average precision of two models trained identically except that one received the measured inverse reduced ion mobility of the precursor as a third precursor feature beside mass and charge, encoded by a sinusoidal float encoder and summed into the same global token. This comparison was carried out on an earlier version of the data corpus, so its absolute values are not comparable with those of the other supplementary figures; only the difference between the two arms is. Supplying the value changed average precision by 0.0003, so the model gains essentially nothing from it; inverse reduced mobility is largely determined by precursor mass and charge, which the model already receives.

#### Supplementary Tables

Supplementary Table 1: **Composition of the training corpus.** Peptidoform counts are taken after the deamidation collapse described in Methods (N[Deamidated] to D and Q[Deamidated] to E), and carbamylation was removed from the corpus. A peptidoform is a sequence with its modifications, a precursor is a peptidoform in one charge state, and a sequence is the peptide with its modifications stripped.

| Split | PSMs | Peptidoforms | Precursors | Sequences |
|---|---|---|---|---|
| Train | 5,434,924 | 1,834,545 | 2,385,801 | 1,589,509 |
| Validation | 679,187 | 228,901 | 298,065 | 224,080 |
| Test | 678,016 | 228,962 | 297,883 | 224,118 |
| Total | 6,792,127 | 2,292,408 | 2,981,749 | 2,037,707 |

Supplementary Table 2: **Residue abundances in the training corpus.** Every occurrence of each residue across the three splits, modified or not. Isoleucine and leucine are listed separately here although the model is evaluated with the two treated as equivalent.

| Residue | Train | Validation | Test | Total | Share |
|---|---|---|---|---|---|
| A | 5,989,764 | 748,296 | 748,380 | 7,486,440 | 7.35% |
| C | 851,721 | 106,282 | 105,505 | 1,063,508 | 1.04% |
| D | 5,331,831 | 665,599 | 667,347 | 6,664,777 | 6.55% |
| E | 7,201,691 | 898,300 | 898,728 | 8,998,719 | 8.84% |
| F | 3,017,437 | 376,745 | 376,178 | 3,770,360 | 3.70% |
| G | 5,158,682 | 642,417 | 644,595 | 6,445,694 | 6.33% |
| H | 2,202,095 | 277,390 | 273,757 | 2,753,242 | 2.70% |
| I | 4,130,490 | 512,676 | 515,577 | 5,158,743 | 5.07% |
| K | 4,522,405 | 564,544 | 564,669 | 5,651,618 | 5.55% |
| L | 7,758,413 | 968,111 | 970,368 | 9,696,892 | 9.52% |
| M | 1,925,333 | 240,235 | 239,376 | 2,404,944 | 2.36% |
| N | 3,127,477 | 391,328 | 391,304 | 3,910,109 | 3.84% |
| P | 4,691,635 | 581,939 | 586,883 | 5,860,457 | 5.76% |
| Q | 3,589,785 | 451,721 | 448,904 | 4,490,410 | 4.41% |
| R | 3,400,763 | 425,248 | 423,847 | 4,249,858 | 4.17% |
| S | 5,901,467 | 736,373 | 737,578 | 7,375,418 | 7.24% |
| T | 4,367,938 | 546,292 | 546,249 | 5,460,479 | 5.36% |
| V | 5,330,026 | 663,478 | 662,847 | 6,656,351 | 6.54% |
| W | 662,430 | 83,083 | 82,455 | 827,968 | 0.81% |
| Y | 2,324,126 | 289,899 | 288,560 | 2,902,585 | 2.85% |
| All residues | 81,485,509 | 10,169,956 | 10,173,107 | 101,828,572 | 100% |

Supplementary Table 3: **Modification abundances in the training corpus.** These five are the complete set of modifications in the model's vocabulary: three residue modifications, of methionine and cysteine, and two N-terminal modifications. The table gives occurrences of each modification across the three splits, the number of spectra whose peptide carries it at least once, and the share of all spectra those represent. Occupancy is the number of occurrences divided by all occurrences of the residue carrying the modification, for instance oxidised methionines out of every methionine; it is not defined for the two N-terminal modifications, which have a single site per peptide.

| Modification | Train | Validation | Test | Total | Spectra | Occupancy |
|---|---|---|---|---|---|---|
| `M[Oxidation]` | 916,476 | 113,499 | 113,357 | 1,143,332 | 1,015,220 | 47.54% |
| `C[Carbamidomethyl]` | 779,787 | 97,188 | 96,482 | 973,457 | 779,061 | 91.53% |
| `C[Cysteinyl]` | 49,691 | 6,202 | 6,136 | 62,029 | 60,812 | 5.83% |
| `[Acetyl]-` | 135,413 | 16,637 | 16,776 | 168,826 | 168,826 | – |
| `[Ammonia-loss]-` | 106,298 | 13,441 | 13,418 | 133,157 | 133,157 | – |
